# Predicting interaction-specific protein–protein interaction perturbations by missense variants with MutPred-PPI

**DOI:** 10.64898/2025.12.20.695738

**Authors:** Ross Stewart, Florent Laval, Georges Coppin, Kerstin Spirohn-Fitzgerald, Maxime Tixhon, Tong Hao, Michael A. Calderwood, Matthew Mort, David N. Cooper, Marc Vidal, Predrag Radivojac

## Abstract

Disruption of protein–protein interactions (PPIs) is a major mechanism of a variant’s deleterious effect. Computational tools are needed to assess such variants at scale, yet existing predictors rarely consider loss of specific interactions, particularly when variants perturb binding interfaces without significantly affecting protein stability. To address this problem, we present MutPred-PPI, a graph attention network that predicts interaction-specific (edgetic) effects of missense variants by operating on AlphaFold 3-based protein complex contact graphs with protein language model embeddings imposed upon nodes. We systematically evaluated our model with stringent group cross-validation as well as benchmark data recently collected within the IGVF Consortium. MutPred-PPI outperformed all baseline methods across all evaluation criteria, achieving an AUC of 0.85 on seen proteins and 0.72 on previously unseen proteins in cross-validation, demonstrating strong generalizability despite scarce training data. To demonstrate biomedical relevance, we applied MutPred-PPI to variants from ClinVar, HGMD, COSMIC, gnomAD, and two *de novo* neurodevelopmental disorder-linked datasets. Disease-associated variants from ClinVar and HGMD showed strong enrichment for both quasi-null and edgetic effects, whereas population variants from gnomAD increasingly preserved interactions with higher allele frequencies. Notably, we observed a strong edgetic disruption signature in highly recurrent cancer variants from both the full COSMIC dataset and a subset of variants from oncogenes. Recurrent tumor suppressor gene variants and autism spectrum disorder-associated variants exhibited moderate quasi-null enrichment, whilst neurodevelopmental disorder-linked variants showed a weak edgetic disruption signature. These results indicate distinct PPI perturbation mechanisms across disease types and show that MutPred-PPI captures functionally relevant molecular effects of pathogenic variants.

## 1 Introduction

Protein–protein interactions (PPIs) govern cellular function, with the human proteome exhibiting up to two million interactions [1–3]. Genetic variants residing in distinct binding interfaces of a protein can have vastly different functional consequences depending upon which PPIs are disrupted [4]. Whereas some variants cause complete loss of function (LoF) and interactions, “edgetic” variants selectively disrupt PPIs while maintaining other interactions [5, 6]. Variants exhibiting these edgetic mechanisms have been implicated in various diseases [5]. For example, cancer-associated mutations in the coiled-coil domain of BRCA1 impair homologous recombination repair by disrupting its interaction with PALB2, while preserving other functions [7, 8]. This interaction-specific nature of variant effects necessitates computational methods capable of predicting interaction perturbations at the resolution of individual protein pairs.

Most variant effect predictors assess overall pathogenicity or other properties without considering specific interaction partners [9, 10]. Methods such as REVEL [11], MutPred2 [12], and AlphaMissense [13] accurately predict variant pathogenicity [14] and have been recommended for clinical use [15], but prediction of interaction-specific effects remains limited. Existing interaction-specific approaches such as SAAMBE-3D [16] and mCSM-PPI2 [17] predict quantitative changes in binding free energy (ΔΔ*G*) induced by mutations in experimental protein complexes, rather than outputting a probabilistic score that indicates PPI perturbation. SWING [18], a recent advance, uses a sliding window approach to model interactions as language, predicting probabilistic interaction-specific perturbations from amino acid sequence alone. However, stringent evaluation on held-out proteins remains necessary to support generalization.

The convergence of recent developments in machine learning and biotechnology platforms is creating new opportunities for accurate prediction of PPI perturbations. AlphaFold 3 [19] mitigates the requirement for experimental structures by providing accurate structural predictions for protein complexes of interest. Protein language models (pLMs), such as ProtT5 [20], generate per-residue embeddings that capture evolutionary and functional information without requiring costly multiple sequence alignments. Furthermore, recent variant interaction profiling through the Impact of Genomic Variation on Function (IGVF) Consortium [21] provides newly generated experimental data for the training and validation of machine learning models.

In this work, we build upon these developments and present MutPred-PPI, a graph attention network (GAT) [22] approach for predicting interaction-specific effects of missense variants. Our method improves upon previous work by: (1) using AlphaFold 3 to generate protein complex structures for interacting protein pairs; (2) exploiting pre-trained pLM embeddings that generalize across protein families; and (3) employing a two-stage training strategy that first learns general stability disruption patterns before fine-tuning on interaction-specific effects. We validate our approach using stringent cross-validation and benchmark schemes that test generalization to unseen proteins and demonstrate superior performance compared to existing methods. Application to disease-associated variants revealed distinct PPI perturbation patterns between disease types, supporting PPI perturbation as a key pathogenic mechanism. Overall, these results provide new insights into molecular mechanisms of genetic disease consequent to mutation.

### 2 Methods

### 2.1 Preprocessing

The end-to-end pipeline of MutPred-PPI is shown in Figure 1. Each data point, which we refer to as a variant-partner pair, is represented as a triplet (*i, v, p*) where *i* is the wild-type interactor protein, *v* specifies the missense variant within *i*, and *p* is a known interaction partner of *i*.

**Figure 1:**
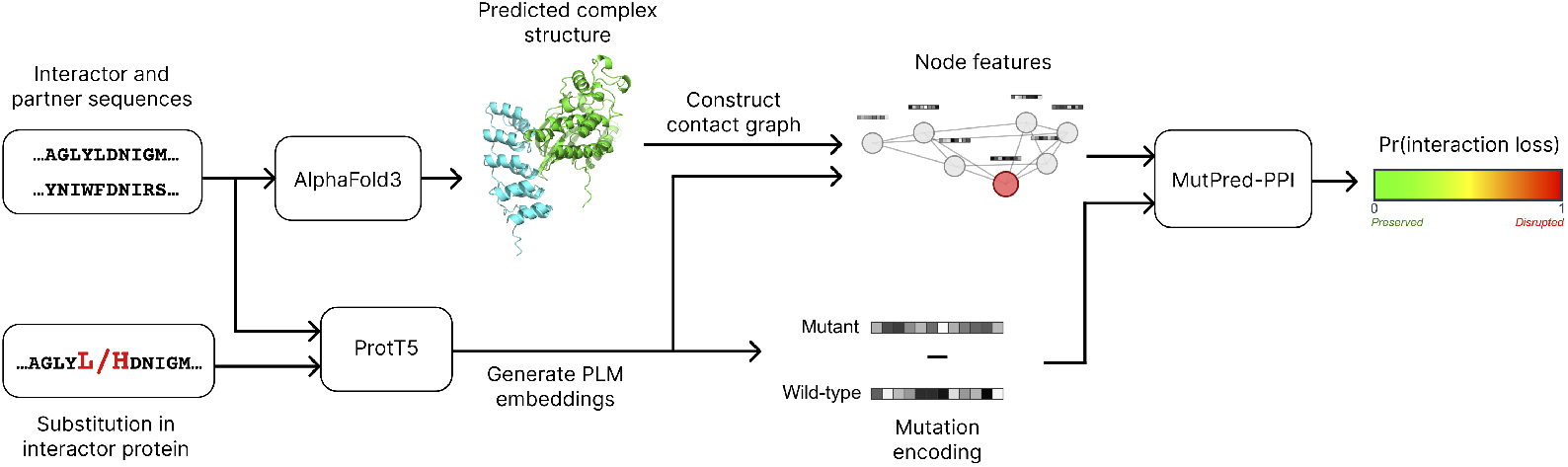
MutPred-PPI end-to-end pipeline. Wild-type sequences of *i* and *p* are input to AlphaFold 3 to generate the complex structure. Wild-type and mutant sequences are input to ProtT5 to generate embeddings, which are passed through MutPred-PPI to give Pr(interaction loss).

First, the amino acid sequences of *i* and *p* are entered into AlphaFold 3 (default parameters) to model the protein complex. Then, we compute the contact graph from the AlphaFold 3 structure, with nodes representing amino acid residues and edges representing contacting residues. We define residues as in contact if any heavy atom distance is ≤ 4.5 °A [23].

Next, we generate pLM embeddings with ProtT5 (ProtT5-XL-UniRef50) [20] for the wild-type sequences of both *i* and *p*, as well as the mutant sequence of *i* (denoted *i*_*v*_). The pLM embeddings of *i*_*v*_ and *p* are then mapped onto the contact graph of the protein complex as node features. We chose to incorporate the mutant embeddings of the interactor protein rather than wild-type embeddings to capture variant-specific context, as computing separate AlphaFold 3 complexes for each variant would have been computationally prohibitive, and AlphaFold models have been shown to be insensitive to structural changes induced by missense variants [24].

Finally, the mutation is encoded as the difference between the mutant and wild-type residue embeddings at the variant position, followed by z-score normalization across variants. We feed both the node features and the mutation encoding into MutPred-PPI to predict the probability of interaction loss.

### 2.2 Model architecture

The model architecture, as shown in Figure 2, consists of three main components: graph attention layers to learn local structural context, a mutation processor to encode mutation effects, and a prediction head for binary classification. Based on the principle that interface mutations are enriched for PPI-perturbing effects [25], our model architecture prioritizes the local structural context at the mutation site where protein–protein contacts are directly affected, rather than global protein features. A detailed mathematical representation of the model architecture is shown in Supplementary Section 1.

**Figure 2:**
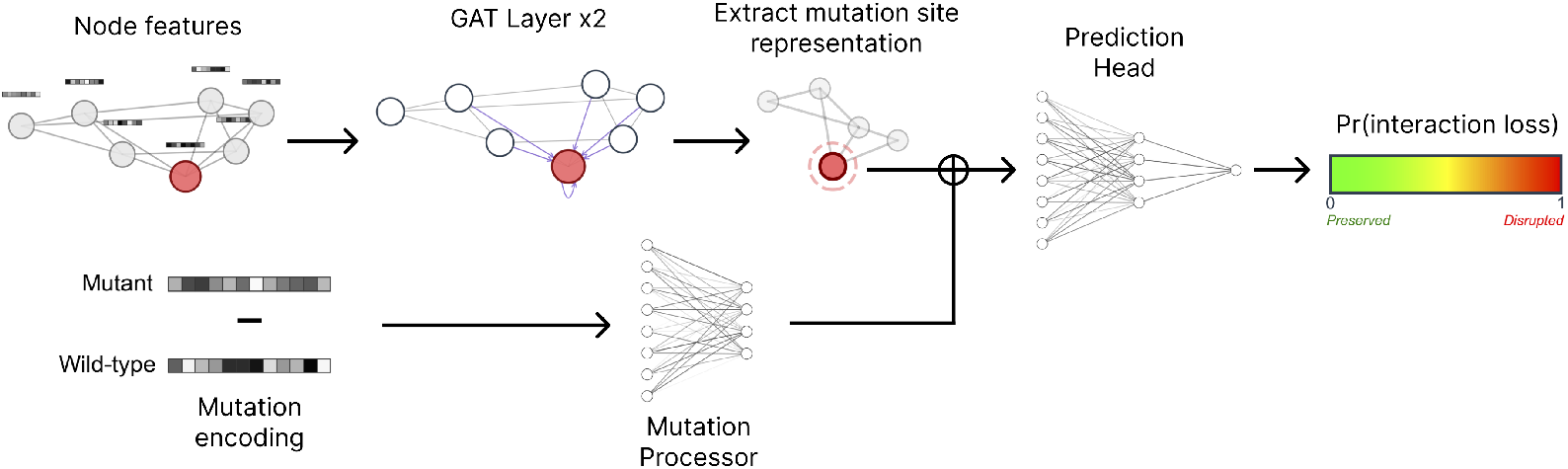
MutPred-PPI model architecture. The node features are propagated through both GAT layers to capture local structural context. The mutation encoding is processed separately, then combined with the node representation at the mutation site for final prediction. Symbol ⊕ denotes concatenation.

### 2.3 Training procedure

We employed a two-stage transfer learning strategy. MutPred-PPI was first pretrained to predict protein stability disruption using experimental structures of monomeric proteins, then fine-tuned for interaction-specific perturbations using AlphaFold 3 complexes. Variants were classified as stability-preserving if |ΔΔ*G*| *<* 0.5 kcal/mol and stability-disrupting if |ΔΔ*G*|≥1.5 kcal/mol [26]. The stability pretraining dataset (176,908 variants) was substantially larger than the available PPI perturbation data (5,894 variant-partner pairs), enabling the model to learn generalizable mutational representations before specializing in edgetic effects. Class weighting was applied to address class imbalance in both pretraining and fine-tuning.

### 2.4 Data

We combined three main datasets for model training and evaluation: PPI perturbation data of variants across Mendelian disorders from Sahni *et al*. [27], population variant PPI perturbation data from Fragoza *et al*. [28], and stability data for pretraining from Blaabjerg *et al*. [29]. For additional performance evaluation, we used in-house benchmark data recently generated through the IGVF Consortium, which we refer to as VarChAMP (Variant Characterization Across the Mendelian Proteome). The VarChAMP dataset was generated through yeast two-hybrid assays using histidine dropout media supplemented with 1 mM 3-Amino-1,2,4-triazole (3-AT), retaining only variants with successfully recovered sequences. Detailed dataset statistics are shown in Table 1. For all datasets, we removed duplicate data points, those with conflicting labels, and those with failed structure generation.

**Table 1:** Dataset statistics for model training and evaluation. Triplets refer to variant-partner combinations.

| Dataset | Proteins | Pairs | Variants | Triplets | Disruptive | Non-disruptive |
| --- | --- | --- | --- | --- | --- | --- |
| *PPI Perturbation Data* |  |  |  |  |  |  |
| Sahni *et al.* (Mendelian) | 537 | 559 | 489 | 1,487 | 532 | 955 |
| Fragoza *et al.* (Population) | 1,868 | 2,252 | 1,904 | 4,515 | 847 | 3,667 |
| VarChAMP (IGVF) | 429 | 436 | 633 | 2,935 | 2,067 | 868 |
| *Combined Training Sets* |  |  |  |  |  |  |
| Sahni, Fragoza | 2,083 | 2,710 | 2,310 | 5,894 | 1,395 | 4,499 |
| Sahni, Fragoza, VarChAMP | 2,367 | 3,118 | 2,918 | 8,827 | 3,466 | 5,361 |
| *Stability Pretraining Data* |  |  |  |  |  |  |
| Blaabjerg *et al.* | 45 | - | 176,908 | - | 112,028 | 64,880 |

### 2.5 Evaluation

To mitigate information leakage commonly arising in paired protein input schemes (due to sequence similarity and shared individual proteins) [30], we first grouped each variant-partner pair by clustering the concatenated sequence of both wild-type proteins. To cluster sequences, we used CD-HIT [31] with a 50% sequence identity threshold. We then evaluated the model through 30 iterations of 10-fold group cross-validation (grouped by sequence cluster) on the training dataset as well as blind-test predictions on the benchmark dataset. To aggregate predictions during cross-validation, we averaged the performance across each fold and iteration, weighted by the number of test data points. For benchmark predictions, we constructed an ensemble of 10 models (one per cross-validation fold), averaging predictions across models. Owing to the heterogeneity between the two PPI perturbation training datasets, we also evaluated the effects of fine-tuning on only the dataset of Mendelian variants versus the combined dataset.

Test predictions were binned into three classes, per Park and Marcotte [30], because protein- and variant-level information leakage can still occur from the pair input scheme. Class one represents predictions with both proteins shared in the training data. Class two represents predictions with one protein shared in the training data. Class three represents predictions with neither protein shared in the training data. Variant-partner pairs are never shared across training and test partitions; however, shared variants may appear in classes one and two with different partner combinations. Class three ensures both proteins and variants are unseen.

We compared MutPred-PPI to SWING [18] (trained on identical cross-validation partitions) and two pretrained proxy models: SAAMBE-3D [16] and MutPred2 [12]. In the official implementation for SWING, a language model is pretrained on training and test data (without labels), and then a discriminator is trained on embeddings generated from that model. To eliminate any information leakage from the test distribution, we evaluated SWING under two scenarios: pretraining on the training and test data (test pretrain), and pretraining on training data only (blind-test). This blind-test training scenario ensured that test data were truly unseen at inference time. To rank SAAMBE-3D predictions, we used the predicted change in binding free energy induced by a mutation (operating on the AlphaFold 3 structure). SAAMBE-3D predictions were binned into test classes based on overlap with its own training data (experimental PDB structures), which differed from our cross-validation splits. MutPred2, a pathogenicity predictor, determined baseline performance because variant pathogenicity can serve as a proxy for interaction loss. However, MutPred2 does not incorporate partner protein information and has constant predictions for a variant across distinct partners (and hence, test classes did not apply). For blind-test evaluation on the benchmark dataset, variants overlapping with the training data of each model were excluded; thus, potential information leakage in classes one and two arose only at the protein level. MutPred2 predictions were not filtered based on variant overlap as MutPred2 was not trained on PPI perturbation data.

### 2.6 Predictions on variant repositories

After fine-tuning on a combined dataset containing the original training data of Mendelian and population variants as well as VarChAMP benchmark data, we applied the resulting ensemble (from cross-validation) to variants from multiple disease and population variant repositories. By aggregating predictions across partners, we predicted edgotypes (i.e., patterns of interaction perturbations induced by a mutation) of variants from ClinVar [32], COSMIC [33], HGMD [34], gnomAD [35], and two *de novo* neurodevelopmental disorder datasets [12, 36]; see Supplementary Section 2 for detailed sources.

First, we gathered variants from each repository alongside their respective partner proteins from BioGRID [37] (physical binding evidence only). Owing to the computational cost of AlphaFold 3, for large datasets, we selected a subset of interactor proteins and up to 10 partner proteins (for each interactor protein) for structure generation and predicted interaction perturbation for all variants in those protein pairs. For variants in the partner protein, we flipped the protein roles (treating the partner as the interactor) before model input. Additionally, we downloaded 126,118 high-confidence precomputed AlphaFold 3 predictions from ProtVar [38] and generated predictions for variants in either protein.

Variants in the partner protein of generated complexes, as well as either protein of precomputed complexes, often had far fewer than 10 partners tested, reducing confidence in edgotype classification. To mitigate this limitation, we only analyzed variants with at least three tested partner proteins when three or more partners existed in BioGRID; otherwise variants were processed normally. Protein pairs with either protein shared in the training dataset were excluded.

Variants were classified as “quasi-wild-type” if all partner interactions were predicted to be unperturbed, “quasi-null” if all were predicted to be perturbed, and “edgetic” if predictions were mixed across partners [27]. Given the class-weighted training objective, we applied a threshold of 0.5 to determine whether an interaction was considered perturbed. Because interaction loss probabilities across partners are not independent, and model outputs may be miscalibrated relative to population frequencies, a probabilistic formulation (e.g., Pr(quasi-null | *i, v*) = ∏_*p*_ Pr(interaction loss | *i, v, p*)) was inappropriate.

### 2.7 Computational efficiency

A single NVIDIA V100 GPU processes approximately 100 variant-partner pairs per minute with pre-computed AlphaFold 3 structures and contact graphs. AlphaFold 3 structures (using MMseqs2 [39] for sequence alignment) and derived contact graphs can take several minutes to generate per protein complex. Structural features should be generated once per protein pair and cached for all variants. ProtT5 embedding generation is the primary computational cost after structure prediction.

## 3 Results

### 3.1 Performance Evaluation

Cross-validation results on the training dataset containing Mendelian and population variants are shown in Figure 3. MutPred-PPI outperformed all compared methods across test classes. SWING (test pretrain) performed well on class one but showed substantial performance degradation on classes two and three. By contrast, MutPred-PPI maintained more stable performance across classes. SAAMBE-3D showed limited performance across classes, and MutPred2 provided a partner-agnostic baseline.

**Figure 3:**
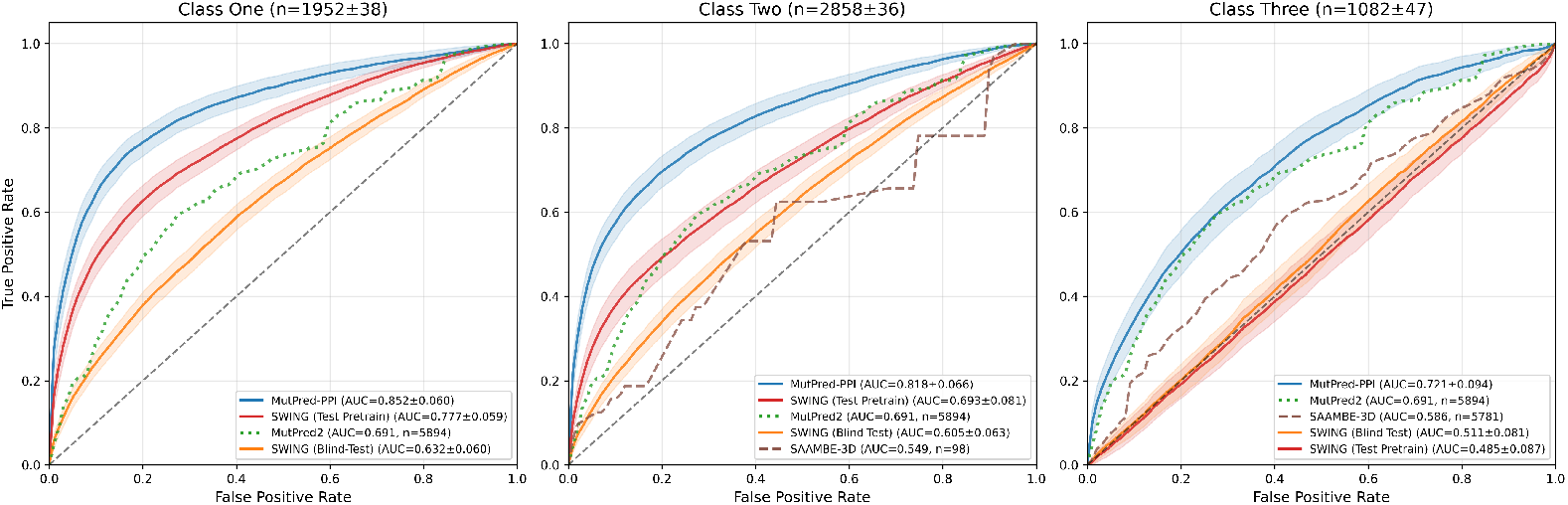
Cross-validation performance on combined Mendelian and population variant training data (*n* = 5,894). Solid lines show mean ROC curves across 30 iterations of 10-fold group cross-validation; shaded regions indicate ± 1 standard error of the mean computed across the 10 folds. Classes one, two, and three represent predictions with both proteins shared, one protein shared, and neither protein shared between training and test data, respectively. MutPred2 predictions are shown constant across test classes (partner-agnostic). Subplot titles show mean test set sizes ±1 standard deviation across the 30 iterations. Proxy methods (dashed and dotted lines) display sample sizes in parentheses. Curves with *n* < 30 were excluded.

The blind-test evaluation results on the VarChAMP benchmark dataset are shown in Figure 4. MutPred-PPI outperformed all compared methods across test classes, with an expected modest performance decline from class one to three. Surprisingly, SWING (test pretrain) often performed worse than SWING (blind-test). MutPred2 achieved comparable performance to MutPred-PPI on class three, and SAAMBE-3D showed minimal predictive signal.

**Figure 4:**
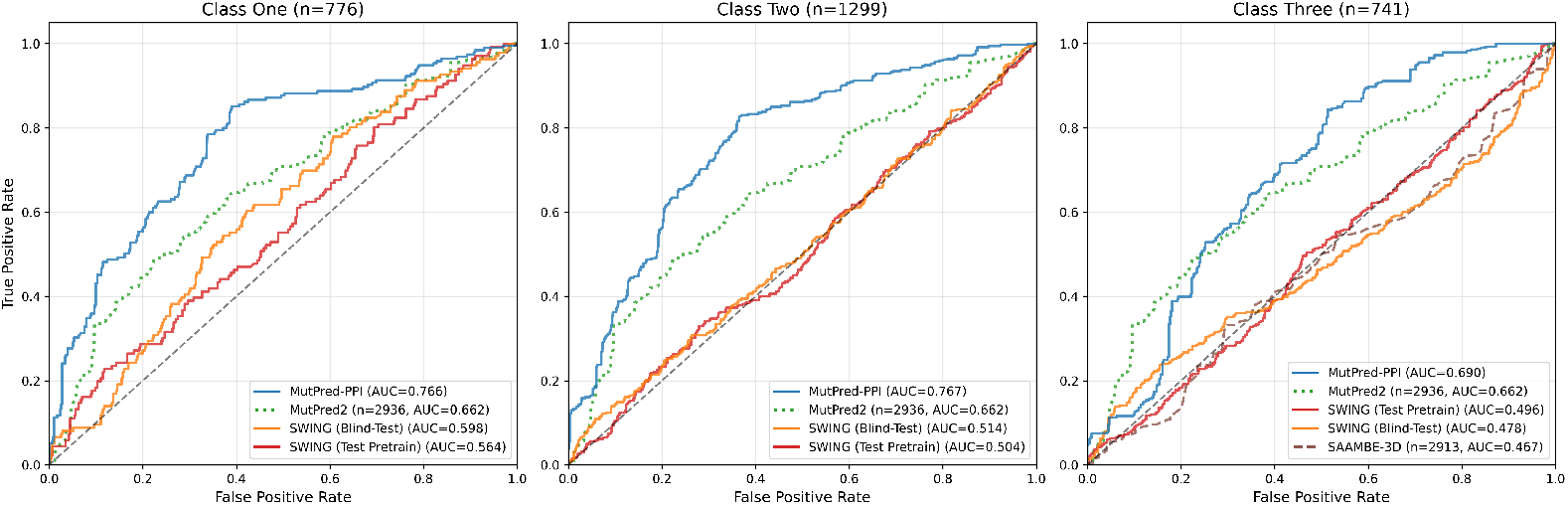
Blind-test ROC curves on the VarChAMP benchmark dataset (*n* = 2,816). Classes one, two, and three represent predictions with both proteins shared, one protein shared, and neither protein shared between training and test data, respectively. MutPred2 predictions are shown constant across test classes (partner-agnostic). Subplot titles show test class sizes. Proxy methods (dashed and dotted lines) display sample sizes in parentheses. Curves with *n* < 30 were excluded.

To compare performance across datasets, we also fine-tuned MutPred-PPI on the Mendelian dataset alone. During cross-validation (Supplementary Figure S1), MutPred-PPI similarly outperformed all baseline methods across test classes. Interestingly, evaluating both MutPred-PPI training configurations on the VarChAMP benchmark dataset (Supplementary Figure S2) showed that including population variants in the fine-tuning dataset reduced class three performance.

### 3.2 Application to disease and population variant repositories

After fine-tuning a final ensemble on the combined training dataset with VarChAMP data (cross-validation performance shown in Supplementary Figure S3), we applied MutPred-PPI to variants from ClinVar, COSMIC, HGMD, gnomAD, and two *de novo* neurodevelopmental disorder datasets. Variant groups were constructed by stratifying each repository: ClinVar by clinical significance; COSMIC by recurrence and gene classification (oncogene vs. tumor suppressor gene, TSG); HGMD as a single group; gnomAD by allele frequency (AF); and two independent *de novo* neurodevelopmental disorder datasets (NDD and ASD) by case/control status. NDD includes cases and controls across four disorders (autism spectrum disorder, intellectual disability, schizophrenia, and epileptic encephalopathy), while ASD contains autism spectrum disorder cases only.

For each variant group, we calculated the enrichment or depletion of quasi-null and edgetic variants relative to the gnomAD population baseline as shown in Figure 5. The enrichment trend *E* was calculated as

**Figure 5:**
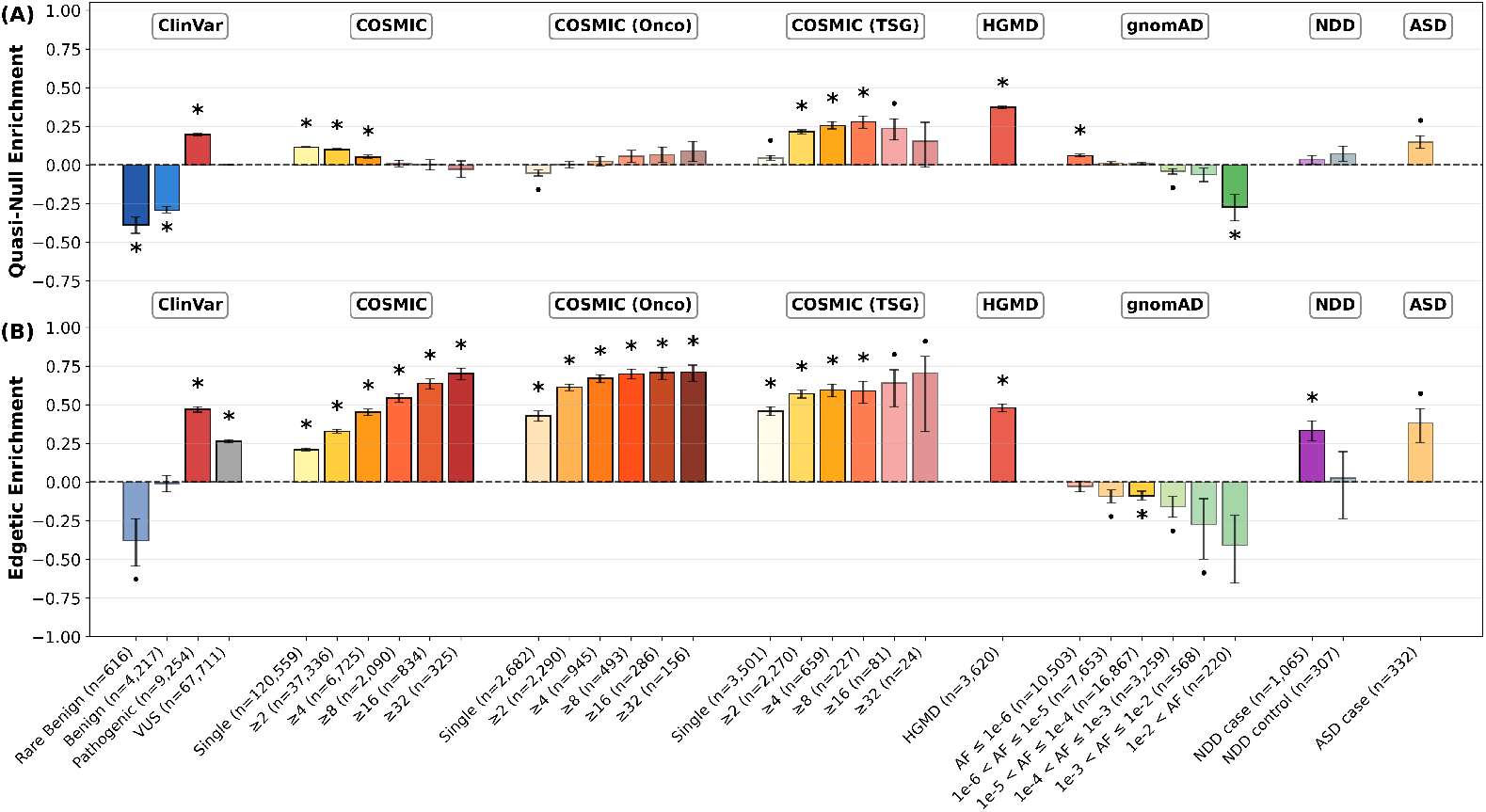
Edgotype enrichment of each variant group with respect to gnomAD across 100,000 bootstrap iterations using all available partners per variant. COSMIC, COSMIC (Onco), and COSMIC (TSG) are binned by recurrence; gnomAD is binned by allele frequency (AF). Rare benign variants are defined as AF ≤ 1%. Error bars indicate 68% confidence intervals. Asterisks indicate statistical significance with Bonferroni correction (*α* = 0.05, *n*tests = 64). Dots indicate significance at the uncorrected *α* = 0.05 level. (A) Quasi-null enrichment. (B) Edgetic enrichment. VUS: variants of uncertain significance.

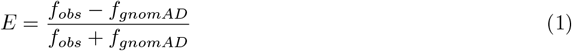

where *f*_*obs*_, *f*_*gnomAD*_ ∈ [0, 1] are the observed rates (relative frequencies) of quasi-null or edgetic variants in the variant group of interest and gnomAD, respectively. The trend is positive if *f*_*obs*_ *> f*_*gnomAD*_ (enrichment) and negative if *f*_*gnomAD*_ *> f*_*obs*_ (depletion). The theoretical range of *E* is [−1, 1], where values approaching 1 indicate strong enrichment and values approaching −1 indicate strong depletion.

Statistical significance was assessed using bootstrap resampling with 100,000 iterations. Enrichments were significant if their sign remained consistent across (1 − *α*_Bonferroni_) of bootstrap samples, with Bonferroni correction (*α* = 0.05, *n*_tests_ = 64) applied across all comparisons.

We observed that variant groups have differential partner coverage (Supplementary Table S1). To account for potential biases this might introduce, we also performed a partner-controlled enrichment analysis by sampling exactly three partners per variant in each bootstrap iteration (Supplementary Figure S4).

#### 3.2.1 ClinVar and HGMD variants

In ClinVar, pathogenic variants show significant enrichment of quasi-null and edgetic effects, while benign variants are depleted for quasi-null effects. Rare benign variants (AF ≤ 1%) exhibit similar depletion. Variants of uncertain significance (VUS) show intermediate PPI perturbation effects between pathogenic and benign, while also being enriched for edgetic effects with respect to the population. HGMD disease-linked variants, similar to ClinVar pathogenic variants, show substantial enrichment for both quasi-null and edgetic effects. In the partner-controlled analysis, these enrichment patterns largely hold, with VUS exhibiting weak enrichment for both quasi-null and edgetic effects, and benign variants showing strong depletion for both effects.

#### 3.2.2 Cancer-associated variants

We analyzed cancer-associated somatic mutations across three gene groups: the full dataset (COSMIC), strictly oncogenes (COSMIC (Onco)), and strictly TSGs (COSMIC (TSG)). In both the full dataset and oncogenes, we observe monotonic trends of increasing edgetic enrichment with recurrence. The full dataset also shows the opposite trend for quasi-null enrichment in statistically significant bins. By contrast, while also exhibiting similar edgetic enrichment trends, TSG variants display a monotonic trend of increasing quasi-null enrichment with recurrence across statistically significant bins, although this trend weakens in the highest recurrence bins ( ≥16 and ≥32) where sample sizes are limited (*n* = 81 and *n* = 24, respectively). These patterns largely persist in the partner-controlled analysis, although edgetic enrichment trends in TSG variants are no longer observed.

#### 3.2.3 Population variants

As allele frequency increases in gnomAD population variants, we observe trends towards depletion of both quasi-null and edgetic effects, exhibiting near-monotonic patterns. In the partner-controlled analysis, these patterns hold although are less extreme.

#### 3.2.4 Neurodevelopmental disorder variants

In the NDD dataset (variants across the four neurodevelopmental disorders), we observe significant edgetic enrichment in case variants but not controls. Case variants from the ASD-specific dataset, although not statistically significant after Bonferroni correction (*n* = 332 variants), exhibit substantial enrichment for quasi-null and particularly edgetic effects. In the partner-controlled analysis, NDD and ASD-specific case variants lose statistically significant edgetic enrichment, and ASD-specific variants show stronger quasi-null enrichment.

#### 3.2.5 Example edgetic variant prediction

To illustrate an edgotype prediction, we examined *BRCA1* p.C61G, a well-characterized pathogenic germline variant (Figure 6). C61 coordinates a zinc ion in the second zinc finger of the BRCA1 RING domain; its substitution to glycine destabilizes local RING structure and abolishes E3 ubiquitin ligase activity [40, 41]. The resulting structural perturbation is primarily confined to the second Zn^2+^-binding loop and does not disrupt overall protein stability [42]. The E2 conjugating enzyme UbcH5c binds directly to the RING zinc-binding loops surrounding C61 [43], and p.C61G reduces BRCA1–BARD1 heterodimerization [44]. MutPred-PPI predicted disruption of the BRCA1–BARD1 interaction as well as interactions with the E2 ubiquitin-conjugating enzymes UBE2D3 (UbcH5c) and UBE2L3 (UbcH7), both of which bind the BRCA1 RING zinc-binding loops [43]. The BRCT-binding partner RBBP4, which interacts with the BRCA1 C-terminal domain [45], was predicted to be preserved.

**Figure 6:**
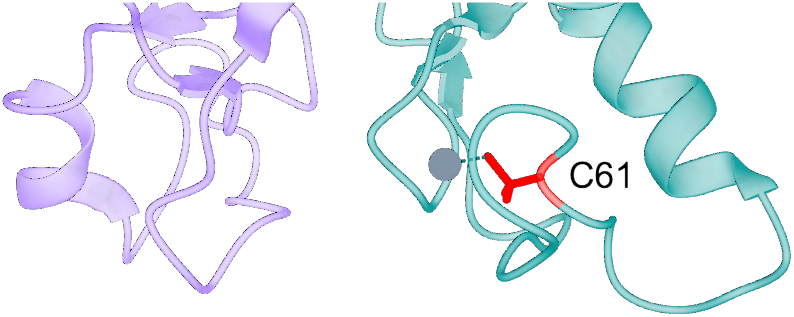
Structure of the BRCA1–BARD1 RING heterodimer (PDB: 1JM7) centered on BRCA1 residue C61. MutPred-PPI (operating on the AlphaFold 3 structure) predicted disruption of the BRCA–BARD1 interaction. Teal: BRCA1; purple: BARD1; red: C61 side chain; gray: zinc ion coordinated by C61.

## 4 Discussion

### MutPred-PPI generalizes to unseen proteins

MutPred-PPI outperformed all compared methods and demonstrated superior generalization to unseen proteins. In cross-validation, the performance of SWING dropped from AUC 0.78 (class one) to 0.49 (class three), while MutPred-PPI maintained more stable performance (0.85 to 0.72). This robustness to unseen proteins illustrates its applicability to proteome-wide variant interpretation.

### Disease variants are enriched for PPI perturbation

Across HGMD and ClinVar, pathogenic variants showed strong enrichment for quasi-null and edgetic effects, while ClinVar benign variants showed strong depletion of quasi-null effects, with depletion patterns persisting in rare benign variants. Disease-associated variants are known to frequently disrupt protein stability [46] and be enriched at PPI interfaces [47]. Our results suggest that pathogenic variants perturb PPIs through LoF and edgetic mechanisms [27], which is consistent with these established principles. The trend of VUS demonstrating an intermediate functional disruption effect between pathogenic and benign variants is also consistent with experimental findings [48].

### Highly recurrent cancer variants are enriched for edgetic mechanisms

The recurrence of variants shows a strong correlation with predicted edgetic effect in cancer variants. This aligns with established cancer biology, as variants with higher recurrence are more likely to be oncogenic drivers [49], and the alteration of PPIs plays a major role in oncogenic signaling [50]. Cancer driver genes are strongly enriched for mutations in PPI interfaces [47, 51], particularly at central network positions [52]. Although MutPred-PPI does not predict gain of interactions, the monotonic trend of increasing edgetic enrichment from single-occurrence to high recurrence observed in both the full dataset of COSMIC variants and the subset of oncogene variants indicates positive selection for network-rewiring mutations in oncogenes. The diminished quasi-null enrichment further suggests that edgetic disruption, rather than protein destabilization, can contribute to gain of function, as demonstrated by recurrent *PIK3CA* helical domain mutations that selectively disrupt inhibitory p85 binding, consequently causing gain of function [53].

By contrast, TSG variants show a monotonic trend of increasing quasi-null enrichment with recurrence in statistically significant bins, while also exhibiting weak edgetic enrichment (edgetic trends are lost in the partner-controlled analysis). This difference reflects distinct selective pressures, where recurrent mutations in tumor suppressors favorably exhibit complete LoF mechanisms, as demonstrated experimentally for *RUNX1* [54]. Additionally, the contrast between enrichment trends of cancer and inherited disease mutations aligns with prior observations that these mutation types exhibit distinct molecular disruption patterns [55].

### Population variants show depletion of PPI perturbation with increasing allele frequency

Among population variants, increasing allele frequencies correlate with benign status due to purifying selection against deleterious mutations [56, 57], and hence PPI-perturbing variants. This depletion trend is observed in both gnomAD predictions and experimentally profiled population variants [28], and contrasts with the enrichment observed in disease-associated variants. Nevertheless, while depleted, common population variants are not necessarily devoid of PPI perturbation mechanisms.

### Neurodevelopmental disorder variants show variable PPI perturbation patterns

NDD case variants show significant edgetic enrichment. However, this differs between analyses, suggesting weak overall enrichment. This weaker signature may reflect that *de novo* missense variants in neurodevelopmental disorders often show more subtle functional effects [58, 59], and that not all *de novo* case variants are causative, unlike ClinVar and HGMD variants with stringent curation criteria. ASD-specific variants exhibit moderate enrichment for both quasi-null and edgetic effects, although the partner-controlled analysis suggests primarily quasi-null enrichment. Larger datasets are needed to more definitively characterize PPI perturbation mechanisms in neurodevelopmental disorders.

### Implications for clinical variant interpretation

The ability of MutPred-PPI to predict interaction-specific effects of mutations has direct implications for clinical variant interpretation. Identifying specific PPIs that are perturbed offers mechanistic insights into disease etiology, revealing disrupted pathways and explaining the molecular basis of pathogenicity. For VUS, the observed intermediate PPI perturbation enrichment between pathogenic and benign variants suggests that MutPred-PPI could aid in variant reclassification, particularly within ACMG/AMP frameworks [60], where interaction-specific predictions provide mechanistic evidence beyond traditional pathogenicity scores, similar to splicing disruption predictions [61]. Our predictions could also prove useful for precision medicine approaches by indicating whether mutating a protein of interest would perturb specific PPIs while leaving others intact.

### Limitations

Our per-variant predictions may underestimate edgetic effects due to limited partner sampling and differential coverage across variant groups. This differential coverage reflects potential BioGRID ascertainment bias [62], and that disease variants are enriched in hub proteins with central network positions [52, 63]. However, our partner-controlled enrichment analysis (Supplementary Figure S4) showed that disease enrichment trends largely hold when controlling for this bias, thereby supporting the biological validity of our findings.

### Future work

The field of predicting interaction-specific effects of mutations remains underdeveloped owing to problem difficulty and the scarcity of high-quality, context-specific experimental data. Continued production of such data remains paramount for advancing computational predictors towards accurate mechanistic models. Further computational development necessitates advances in predictor accuracy, prediction of interaction gain, extensions to other macromolecules and ligands, and coverage of additional variant types. The consequences of alternative splicing on PPI perturbations also remain a challenging prediction task [64] and could be addressed through a modified variant effect prediction framework. As these methods mature, they will become increasingly valuable for understanding how genetic variation causes disease through interactome-level effects.

## Acknowledgements

We thank the VarChAMP group from the IGVF Consortium for providing access to the benchmark dataset. We acknowledge the use of AlphaFold 3 (Google DeepMind) and the Catalogue of Somatic Mutations in Cancer (COSMIC, Wellcome Sanger Institute) under academic licenses. This work was supported by the NIH awards U01HG012022 (P.R.), R01GM145937 (P.R.), and UM1HG011989 (M.V.).

## Competing Interests

The authors declare the following competing interests. D.N.C. and M.M. acknowledge Qiagen Inc. for their financial support through a License Agreement with Cardiff University. The remaining authors declare no competing interests.

## Code and Data Availability

Source code and trained models are available at https://github.com/rosstewart/MutPred-PPI. The processed datasets used for training, evaluation, and predictions are available at https://doi.org/10.5281/zenodo.17645488. Provided AlphaFold 3 predictions are subject to AlphaFold Server Output Terms of Use [19]. VarChAMP data were unpublished at the time of submission and hence were excluded. We were unable to deposit COSMIC and HGMD variant-partner interactions due to licensing restrictions; these should be obtained directly from their respective sources.

## Notes

https://github.com/rosstewart/MutPred-PPI

https://doi.org/10.5281/zenodo.17645488

### Summary of Updates

New results figure; minor changes throughout the manuscript.

