## Supplementary Information for "Predicting interaction-specific protein–protein interaction perturbations by missense variants with MutPred-PPI"

**2 Variant repositories data sources** **3**

**3 Supplementary figures** **5**

### 1 Detailed model architecture

#### 1.1 Input representation

The input to the model consists of: a node feature matrix $\mathbf{X} \in \mathbb{R}^{n \times d}$, where $n$ is the length of the concatenated sequences of proteins $i$ and $p$, and $d = 1024$ represents the embedding dimension of ProtT5. The adjacency matrix $\mathbf{A} \in \{0, 1\}^{n \times n}$ encodes residue contacts. For each variant, let $k \in \{1, \ldots, |i|\}$ be the residue index of the mutation in the interactor protein. We compute the mutation encoding $\widetilde{\Delta \mathbf{x}}_k \in \mathbb{R}^d$ as the normalized difference between the mutant and wild-type embeddings at position $k$.

#### 1.2 Graph attention layers

The node features are processed through two GAT layers. Each GAT layer computes updated node representations using multi-head attention:

$$\mathbf{h}_i^{(l+1)} = \sigma \left( \frac{1}{K} \sum_{k=1}^{K} \sum_{j \in \mathcal{N}_i} \alpha_{ij}^{k,(l)} \mathbf{W}^{k,(l)} \mathbf{h}_j^{(l)} \right) \tag{1}$$

where $\mathbf{h}_i^{(l)}$ is the representation of node $i$ at layer $l$, $\mathbf{W}^{k,(l)}$ is the learnable weight matrix for attention head $k$ at layer $l$, $K$ is the number of attention heads, $\mathcal{N}_i$ represents the one-hop neighborhood of node $i$ defined by the adjacency matrix, and $\alpha_{ij}^{k,(l)}$ are the attention coefficients for head $k$ in layer $l$. The attention coefficients are computed as follows:

$$\alpha_{ij}^{k,(l)} = \frac{\exp\left(\text{LeakyReLU}\left(\mathbf{a}^{k,(l)T}[\mathbf{W}^{k,(l)}\mathbf{h}_i^{(l)} \parallel \mathbf{W}^{k,(l)}\mathbf{h}_j^{(l)}]\right)\right)}{\sum_{m\in\mathcal{N}_i}\exp\left(\text{LeakyReLU}\left(\mathbf{a}^{k,(l)T}[\mathbf{W}^{k,(l)}\mathbf{h}_i^{(l)} \parallel \mathbf{W}^{k,(l)}\mathbf{h}_m^{(l)}]\right)\right)} \tag{2}$$

where $\parallel$ denotes concatenation and $\mathbf{a}^{k,(l)}$ is a learnable attention vector. Specifically, our architecture uses the following:

$$\mathbf{H}^{(1)} = \text{ReLU}\left(\text{GAT}_1(\mathbf{X}, \mathbf{A}; \theta_1)\right) \in \mathbb{R}^{n\times 4h} \tag{3}$$

$$\mathbf{H}^{(2)} = \text{ReLU}\left(\text{GAT}_2(\mathbf{H}^{(1)}, \mathbf{A}; \theta_2)\right) \in \mathbb{R}^{n\times h/2} \tag{4}$$

where the first layer uses $K = 4$ attention heads with hidden dimension $h = 256$ (concatenated), and the second layer uses a single head.

#### 1.3 Mutation processing

The normalized mutation encoding is processed through a dedicated neural network:

$$\mathbf{m} = f_{\text{mut}}(\tilde{\Delta\mathbf{x}}_k; \phi) = \text{MLP}_\phi(\tilde{\Delta\mathbf{x}}_k) \in \mathbb{R}^{h/2} \tag{5}$$

where $f_{\text{mut}}$ is a multilayer perceptron (MLP) with one hidden layer, ReLU activation, and dropout:

$$\mathbf{z}_1 = \text{Dropout}_{0.1}(\text{ReLU}(\mathbf{W}_1\tilde{\Delta\mathbf{x}}_k + \mathbf{b}_1)) \tag{6}$$

$$\mathbf{m} = \mathbf{W}_2\mathbf{z}_1 + \mathbf{b}_2 \tag{7}$$

#### 1.4 Prediction head

The final prediction combines the local structural context at the mutation site with the processed mutation encoding:

$$\mathbf{c} = [\mathbf{h}_k^{(2)} \parallel \mathbf{m}] \in \mathbb{R}^h \tag{8}$$

where $\mathbf{h}_k^{(2)}$ is the learned representation of the mutation site residue after two GAT layers. The probability of interaction loss is computed as:

$$\Pr(\text{interaction loss} \mid i, v, p) = \sigma\left(f_{\text{pred}}(\mathbf{c}; \psi)\right) \tag{9}$$

where $f_{\text{pred}}$ is an MLP with two hidden layers:

$$\mathbf{o}_1 = \text{Dropout}_{0.1}(\text{ReLU}(\mathbf{W}_3\mathbf{c} + \mathbf{b}_3)) \in \mathbb{R}^{h/2} \tag{10}$$

$$\mathbf{o}_2 = \text{Dropout}_{0.1}(\text{ReLU}(\mathbf{W}_4\mathbf{o}_1 + \mathbf{b}_4)) \in \mathbb{R}^{h/8} \tag{11}$$

$$\text{logit} = \mathbf{W}_5\mathbf{o}_2 + \mathbf{b}_5 \in \mathbb{R} \tag{12}$$

The complete model can be summarized as follows:

$$\Pr(\text{interaction loss} \mid i, v, p) = \sigma\left(f_{\text{pred}}\left([\mathbf{h}_k^{(2)} \parallel f_{\text{mut}}(\tilde{\Delta\mathbf{x}}_k)]\right)\right) \tag{13}$$

where all parameters $\{\theta_1, \theta_2, \phi, \psi\}$ are learned through binary cross-entropy loss minimization.

#### 2 Variant repositories data sources

To construct an interactome network, we downloaded all interacting human protein pairs with physical binding evidence from BioGRID Release 4.4.244. We then gathered variants from multiple repositories for analysis. From ClinVar [1] (January 2, 2025 release), we collected pathogenic (P/LP) and benign (B/LB) missense variants with at least one review star, as well as all missense variants of uncertain significance (VUS). All cancer-associated missense variants, recurrence counts, and gene classifications (oncogene vs. tumor suppressor gene, TSG) were downloaded from COSMIC [2] v101 (accessed under academic research terms). To define the oncogene and TSG sets, we retained only genes assigned to a single class. Disease-linked missense variants from HGMD [3] Professional 2025 were downloaded and mapped to amino acid substitutions using ANNOVAR [4], retaining only "DM" variants while excluding "DM?" variants. Population variants were obtained from gnomAD [5] v4.1.0 across all chromosomes and mapped to amino acid substitutions using ANNOVAR. Allele frequencies (AFs) were assigned using the GroupMax flag. Finally, for the two *de novo* variant datasets linked to neurodevelopmental disorders: one (NDD dataset [6]) contained case and control variants from autism spectrum disorder (ASD), intellectual disability, schizophrenia, and epileptic encephalopathy, and the other (ASD-specific dataset [7]) contained case variants specifically associated with ASD. Detailed dataset statistics are available in Supplementary Table S1. For each variant group, we report the number of unique proteins, protein pairs, variants, triplets (variant-partner combinations), and mean partners (the mean number of partners tested per variant).

Table S1: Dataset statistics for variant repositories

| **Dataset** | **Proteins** | **Pairs** | **Variants** | **Triplets** | **Mean Partners** |
|---|---|---|---|---|---|
| *ClinVar* | | | | | |
| Rare Benign | 630 | 559 | 616 | 2,054 | 3.3 |
| Benign | 1,964 | 2,302 | 4,217 | 20,057 | 4.8 |
| Pathogenic | 1,409 | 1,572 | 9,254 | 72,143 | 7.8 |
| VUS | 2,609 | 3,604 | 67,711 | 313,193 | 4.6 |
| *COSMIC* | | | | | |
| Single-occurrence | 2,447 | 3,136 | 120,559 | 431,607 | 3.6 |
| Recurrence $\geq$ 2 | 2,425 | 3,717 | 37,336 | 155,962 | 4.2 |
| Recurrence $\geq$ 4 | 2,091 | 2,837 | 6,725 | 35,895 | 5.3 |
| Recurrence $\geq$ 8 | 1,310 | 1,436 | 2,090 | 13,678 | 6.5 |
| Recurrence $\geq$ 16 | 700 | 695 | 834 | 6,304 | 7.6 |
| Recurrence $\geq$ 32 | 354 | 355 | 325 | 2,704 | 8.3 |
| *COSMIC (Oncogenes)* | | | | | |
| Single-occurrence | 110 | 220 | 2,682 | 14,079 | 5.2 |
| Recurrence $\geq$ 2 | 671 | 1,431 | 2,290 | 12,873 | 5.6 |
| Recurrence $\geq$ 4 | 630 | 1,217 | 945 | 5,416 | 5.7 |
| Recurrence $\geq$ 8 | 576 | 980 | 493 | 2,923 | 5.9 |
| Recurrence $\geq$ 16 | 374 | 599 | 286 | 1,695 | 5.9 |
| Recurrence $\geq$ 32 | 238 | 343 | 156 | 927 | 5.9 |
| *COSMIC (Tumor Suppressor Genes)* | | | | | |
| Single-occurrence | 143 | 286 | 3,501 | 21,441 | 6.1 |
| Recurrence $\geq$ 2 | 943 | 1,738 | 2,270 | 20,835 | 9.2 |
| Recurrence $\geq$ 4 | 902 | 1,442 | 659 | 7,632 | 11.6 |
| Recurrence $\geq$ 8 | 630 | 906 | 227 | 2,947 | 13.0 |
| Recurrence $\geq$ 16 | 269 | 381 | 81 | 1,017 | 12.6 |
| Recurrence $\geq$ 32 | 98 | 132 | 24 | 282 | 11.8 |
| *HGMD* | | | | | |
| All | 1,391 | 1,545 | 3,620 | 21,458 | 5.9 |
| *gnomAD* | | | | | |
| All | 1,073 | 1,118 | 249,316 | 678,231 | 2.7 |
| AF $\leq$ 1e-6 | 431 | 350 | 10,503 | 22,765 | 2.2 |
| 1e-6 $<$ AF $\leq$ 1e-5 | 456 | 371 | 7,653 | 16,064 | 2.1 |
| 1e-5 $<$ AF $\leq$ 1e-4 | 464 | 380 | 16,867 | 34,898 | 2.1 |
| 1e-4 $<$ AF $\leq$ 1e-3 | 404 | 316 | 3,259 | 6,548 | 2.0 |
| 1e-3 $<$ AF $\leq$ 1e-2 | 294 | 215 | 568 | 1,072 | 1.9 |
| 1e-2 $<$ AF | 183 | 131 | 220 | 377 | 1.7 |
| *Neurodevelopmental Disorders* | | | | | |
| Case | 3,359 | 5,694 | 1,065 | 8,338 | 7.8 |
| Control | 2,157 | 2,744 | 307 | 3,045 | 9.9 |
| *Autism Spectrum Disorder* | | | | | |
| Case | 558 | 591 | 332 | 5,831 | 17.6 |

### 3 Supplementary figures

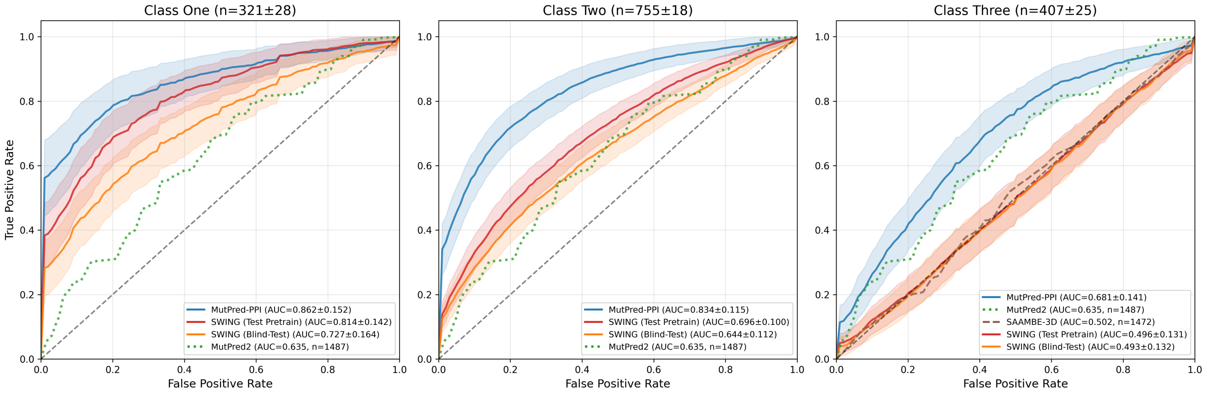

Figure S1: Cross-validation performance on Mendelian variant training data ($n$ = 1,487). Solid lines show mean ROC curves across 30 iterations of 10-fold group cross-validation; shaded regions indicate $\pm 1$ standard error of the mean computed across the 10 folds. Classes one, two, and three represent predictions with both proteins shared, one protein shared, and neither protein shared between training and test data, respectively. MutPred2 predictions are shown constant across test classes (partner-agnostic). Subplot titles show mean test set sizes $\pm 1$ standard deviation across the 30 iterations. Proxy methods (dashed and dotted lines) display sample sizes in parentheses. Curves with $n < 30$ were excluded.

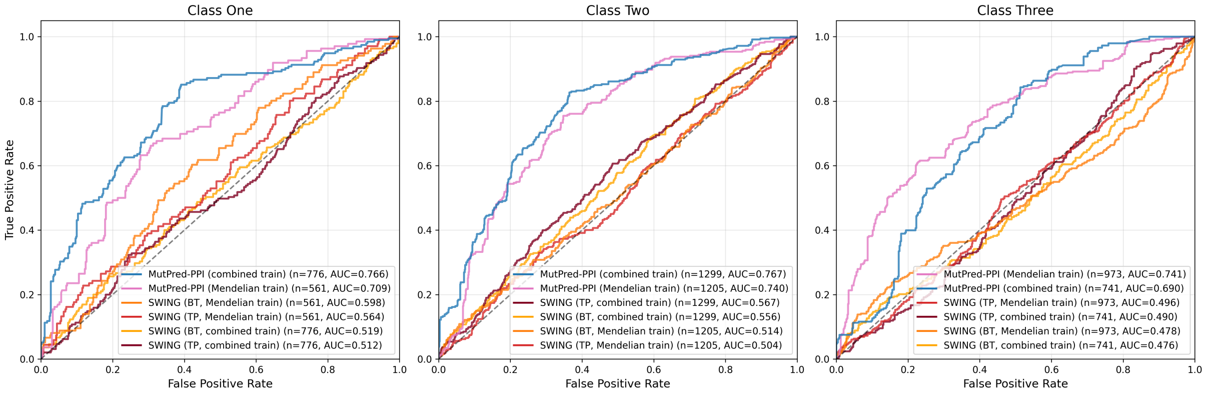

(a) VarChAMP dataset

Figure S2: Blind-test ROC curves on the VarChAMP dataset ($n$ = 2,739–2,816) comparing training on both Mendelian and population variants vs. Mendelian variants only. Classes one, two, and three represent predictions with both proteins shared, one protein shared, and neither protein shared between training and test data, respectively. TP: test pretrain, BT: blind-test.

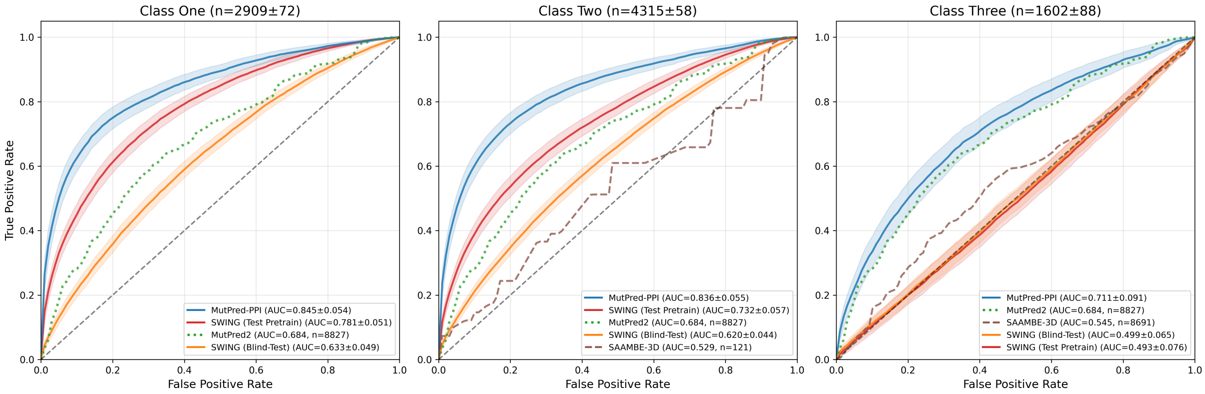

Figure S3: ROC curves of cross-validation predictions on the combined Mendelian, population, and VarChAMP datasets ($n$ = 8,827). Solid lines show mean ROC curves across 30 iterations of 10-fold group cross-validation; shaded regions indicate $\pm 1$ standard error of the mean computed across the 10 folds. Classes one, two, and three represent predictions with both proteins shared, one protein shared, and neither protein shared between training and test data, respectively. MutPred2 predictions are shown constant across test classes (partner-agnostic). Subplot titles show mean test set sizes $\pm 1$ standard deviation across the 30 iterations. Proxy methods (dashed and dotted lines) display sample sizes in parentheses. Curves with $n < 30$ were excluded.

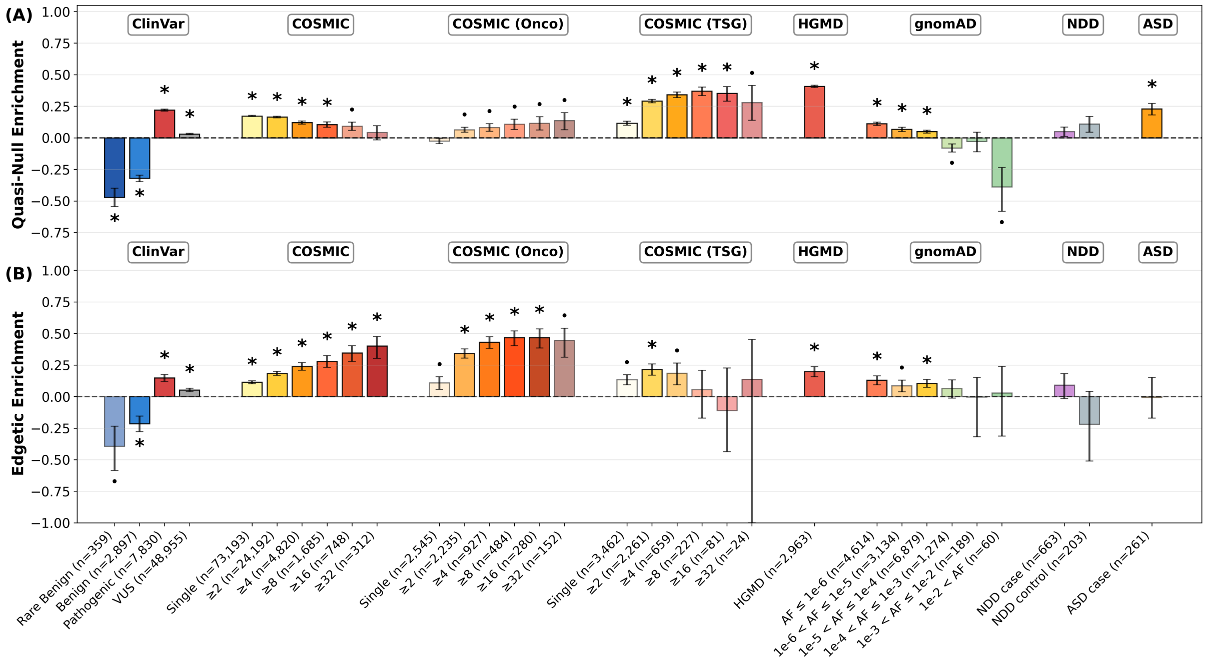

Figure S4: Partner-controlled edgotype enrichment with three partners sampled per variant. For each of 100,000 bootstrap iterations, three partner proteins were randomly sampled for each variant; variants with fewer than three partners were excluded. COSMIC, COSMIC (Onco), and COSMIC (TSG) are binned by recurrence; gnomAD is binned by allele frequency (AF). Rare benign variants are defined as AF $\leq$ 1%. Error bars indicate 68% confidence intervals. Asterisks indicate statistical significance with Bonferroni correction ($\alpha = 0.05$, $n_{\text{tests}} = 64$). Dots indicate significance at the uncorrected $\alpha = 0.05$ level. (A) Quasi-null enrichment. (B) Edgetic enrichment. VUS: variants of uncertain significance.
